# Native experience modulates neural timing: plasticity in language prediction hierarchies

**DOI:** 10.1101/245696

**Authors:** Francesco Giannelli, Nicola Molinaro

**Affiliations:** University of Barcelona, Department of Cognition, Development and Educational Psychology; BCBL, Basque center on Cognition Brain and Language, Donostia-San Sebastian, Spain; Ikerbasque, Basque Foundation for Science, Bilbao, Spain

**Keywords:** language prediction, bilingualism, processing hierarchy, word anticipation, grammatical gender, ERP

## Abstract

We investigated how native language experience shapes prediction mechanisms. Two groups of bilinguals (either Spanish or Basque natives) performed a word matching task (WMT) and a picture matching task (PMT). They indicated whether the stimuli they perceived matched with the noun they heard. Spanish noun endings were either diagnostic of the gender (transparent), or ambiguous (opaque). ERPs were time-locked to the gender-marked determiner preceding the predicted noun. In the WMT both groups showed a negative (~340 ms) effect. Basque natives displayed an earlier effect (~150 ms) for determiners preceding transparent nouns. In the PMT both groups showed an early effect (~160 ms) for determiners preceding opaque nouns. Transparent nouns' determiners elicited prediction at ~330 ms in Spanish natives, but at ~460 ms in Basque natives. It is concluded that bilinguals rely on the features of the L1 for predicting in the L2: native experience moulds prediction. Linguistic prediction is hierarchical, with different operation levels communicating at an interface stage.

## Introduction

Language processing robustly relies on predictive processing mechanisms (Bar, 2007; Clark, 2013; Federmeier, 2007; Friston, 2010; Levy, 2008; Pickering & Garrod, 2013; but see Jackendoff, 2002; Morris, 2006 for a different point of view).

General world knowledge and contextual constraints constantly influence the speakers’ processing system. Top-down operations not only facilitate the incorporation of new inputs into the preceding structure after they have been observed (*integration*), but also pre-activate upcoming inputs before they are even perceived (*prediction*; e.g., Altmann & Mirkovic, 2009; Jaeger & Snider, 2013; Kuperber & Jaeger, 2016; Levy, 2008; MacDonald, 2013).

Anticipatory behaviour in the native language has been attested by a great number of studies. Experiments using the visual word paradigm have shown that when given a constraining context, participants tend to move their eyes towards objects or images before they are explicitly revealed by the input (Altmann & Kamide, 1999; Kamide, Altmann & Haywood, 2003). Research using Event-Related Potentials (ERPs) manipulated context sentences in order to measure what happens in the brain right before participants perceive an unexpected lexical item (in comparison with an expected one), and found replicable electrophysiological correlates of lexical prediction, namely a more negative N400 effect (DeLong, Urbach & Kutas, 2005; Wicha, Moreno & Kutas, 2003). Also, it has been found that oscillatory EEG activity around 4 Hz reflects the degree to which a certain item can be predicted (Molinaro, Barraza, & Carreiras, 2013). When they listen or read, native speakers are able to predict the phonological (DeLong et al., 2005), orthographic (Hawelka et al., 2015), syntactic (Levy, 2008; Dikker et al., 2010) and semantic (Altmann & Kamide, 1999; Kutas & Hillyard, 1984) representation of the input coming next. In the case of mismatch between the predicted and the actual input, language users take advantage of the *prediction error* to adjust future predictions, with the result of speeding up processing, enhancing communication and boosting learning (Chang, Dell, & Bock, 2006; Federmeier, 2007; Pickering & Garrod, 2013).

However, even if predictive sentence processing in the first language (L1) has been widely studied, a comprehensive picture of this phenomenon is still far from being complete.

Much is known about what native speakers are able to predict, but much less is known about when and how they do it. Language processing is extremely fast, and prediction is (by definition) even faster, therefore anticipation processes (and their relative time course) are very hard to grasp: what are the mechanisms supporting language prediction, and when do they operate?

In our opinion, research on second language (L2) anticipation mechanisms is a valid approach to this issue and it is fundamental for comprehending the general mechanisms underlying language prediction in general. When processing a message in an L2, most speakers are not fast enough to keep pace with the speed with which a sentence unfolds, and their comprehension is often impaired (Hahne, 2001;Hahne & Friederici, 2001). As a consequence, even at good levels of proficiency, the ability to anticipate might be reduced, or delayed or it might be different in quality compared to L1 processing. These differences can shed light on the type of representation that the human brain predicts. The present study examines the anticipation processes that are at play during L2 lexical pre-activation when L2 proficiency is very high (thus comparable with L1 prediction processes).

Through analysis of bilingual speakers, this experiment not only aims at finding the differences between L1 and L2 in anticipation processing, but also has the goal of explaining these differences (if any) within a more general and unified theory of the prediction mechanisms underlying language processing.

Initial research on bilingual prediction has shown that non-native speakers differ from natives in terms of quantity and quality of exposure to the L2: this leads to reduced proficiency and impaired prediction mechanisms (Kaan, 2014). Nonetheless, the efficiency of the prediction mechanism would directly depend on the proficiency in each language, therefore it seems reasonable to assume that when L2 proficiency is high enough, L2 prediction mechanism are very much like the ones used in the L1.

In order to shed light on this matter, Martin et al. (2013) investigated lexical prediction, comparing a group of late high proficient Spanish-English bilinguals and a group of English monolinguals. They used an ERP paradigm similar to DeLong et al. (2005) to study the differences between L1 and L2 in the way semantic processing is modulated by lexical pre-activation. Participants read English highly constraining sentences like *He was very tired so he sat on…* By the time they read the sentence, subjects had already predicted the exact noun that would have come next. Nonetheless, sentences could be followed by either an expected or an unexpected (but correct) noun phrase (noun preceded by its article) such as *a chair* or *an armchair*. Crucially, the experiment utilized the phonological property of English according to which the indefinite article “*a*” changes to “*an*” if the noun that follows it starts with vowel, and took advantage of this rule to detect prediction.

In monolinguals, the ERPs recorded on the unexpected article (preceding unexpected nouns) elicited more negative N400 amplitudes than the expected article, in line with DeLong et al. (2005). In contrast, highly proficient bilinguals did not show any prediction effect in the L2. Based on this result, the authors concluded that L2 users do not predict to the same extent native speakers do.

This seemed a robust result (see also Ito et al., in press) but did not take in to account either the typological differences between the languages, or the impact that cross-linguistic similarities may have on anticipation processes.

A follow up study (Foucart et al., 2014) compared Spanish monolinguals, Spanish-Catalan early bilinguals and French-Spanish late bilinguals. The experiment shadowed Wicha et al. (2004): participants read sentences in Spanish where the context was manipulated in a way that the critical noun and its preceding article could be expected or unexpected. Critically, the gender of the expected noun was the opposite of the unexpected ones, and their corresponding articles agreed with them (e.g., *The pirate had the secret map, but he never found the* [masc] *treasure* [masc] / *the* [fem] *cave* [fem]).

ERPs recorded on the unexpected article elicited a more negative N400 effect than the expected articles. Interestingly, the same N400 effect was found in all three groups of participants, regardless of the L1 they spoke, their L2 proficiency or their age of L2 acquisition.

In contrast with the previous experiment, in this study the conclusion was that bilinguals can predict upcoming words in the same way monolinguals do.

The main difference between the two experiments was the linguistic feature manipulated to detect prediction mechanisms. While in the first experiment the phonotactic agreement feature (“*a*”, “*an*”) exists in English but not in Spanish, the gender agreement feature between article and noun is present in Spanish, French and Catalan.

This final distinction opened up several central questions: do typological language differences have an impact on L2 prediction processes? Can bilinguals predict the features of the L2 even though those features are not part of their L1? Again, is it just a matter of proficiency, or does the L1 have an irrecoverable impact on L2 anticipation mechanisms?

### Hierarchical levels of representation in bilingual prediction

We aimed to disentangle these matters in a research project focused on studying prediction by testing two groups of very early bilinguals who spoke two typologically different languages. In a recent EEG experiment (Molinaro et al., 2016; under review) we tested Basque (L1)-Spanish (L2) and Spanish (L1)-Basque (L2) speakers; all of them were highly proficient in both languages but they were different in terms of age of acquisition, with the L2 acquired at the age of three years old.

As in the previous experiments (Foucart et al., 2014; Wicha et al., 2004), subjects read sentences (in Spanish) in which a constraining context was followed by an expected or an unexpected noun phrase, where the latter was formed by a gender marked article and a gender marked noun. Here again, expected and unexpected nouns differed in gender, so that it was possible to observe anticipation effects on the article. In comparison with the previous studies, however, there was a significant difference, as we also considered the hierarchical level at which the gender information is available for the predicted noun.

The Basque language does not have grammatical gender, and its morphological regularities are based on post-nominal suffixes (Laka, 1996; Rijk, 2008). On the contrary, in Spanish, grammatical gender is a feature that is assigned to any inanimate noun, but the noun ending is not always diagnostic of the gender as there is a substantial amount of irregularity. Two thirds of Spanish nouns are gender-transparent (ending in-a for feminine and in-o for masculine nouns), but the remaining nouns are gender-opaque (ending in a consonant, or in a different non-diagnostic vowel) (R.A.E., 2010).

It has been hypothesized that grammatical gender is extracted on the basis of two different routes: one based on the lexical properties (abstract features) of the noun stored in the mental lexicon, and one deriving the gender features from the formal properties of the noun. When the lexical route is not available, language users rely on the formal route to extract gender (Gollan and Frost, 2001).

According to this theory, as Spanish opaque nouns do not have formal cues for gender retrieval, grammatical gender necessarily has to be derived from the lexical route. In contrast, transparent nouns present reliable formal cues (e.g. gender-related noun endings), therefore both the lexical route and the form-based route are utilizable for gender extraction (see Caffarra et al., 2014, for supporting data).

In Molinaro et al. (2016; under review), not only did expected and unexpected nouns differ in gender, but transparency was also taken into account. Participants read sentences that could end with a transparent or an opaque noun, such as *Acabo de salir de la casa y no recuerdo si he cerrado la puerta/el armario”* (“I just got out of the house, and I don’t remember whether I closed the [fem] door [fem, transp] / the [masc] wardrobe [masc,transp]”) or *Prefiero que el te esté muy dulce, puedes pasarme el azúcar / la miel por favor?* (“I prefer my tea very sweet, would you pass me the [masc] sugar [masc, opaque] / the [fem] honey [fem, opaque] ?”).

ERPs recorded on the unexpected determiner (in comparison with the expected) revealed an N400 effect in both groups, independently of the transparency of the predicted lexical element. This result not only replicated Foucart et al. (2014), and demonstrated that highly proficient bilinguals can predict the way monolinguals do, but did so by using a feature, namely gender, which is not available in the L2 of the participants.

In addition, the ERP analysis presented an even more striking outcome. On the unexpected determiners that preceded transparent words, Basque natives showed a P200 effect (starting ~100 ms before the observed N400s), a component classically thought to reflect visual-attention processes (Luck & Hillyard, 1994; Liu et al., 2013; Molinaro et al., 2013; Su et al., 2016).

In order to explain these results we formulated a multifaceted hypothesis. Spanish natives do not rely on formal cues (i.e. *a/o* noun ending) to compute agreement dependencies involving grammatical gender (since ~1/3 of the nouns are gender opaque) (Caffarra & Barber, 2015), but rely on the lexical information stored in the mental lexicon to predict the gender of the word that is coming next, without taking into account the noun transparency, as reflected by the N400 effect.

On the other hand, Basque natives rely more on sublexical, word-form related analysis for the gender prediction of transparent words, as reflected in the P200. They perform the same kind of prediction for opaque words and necessary fail because they have no formal cues to base their prediction on, but they can still switch to the lexical route in order to predict the gender. The reason for this processing difference would reside in the fact that in Basque, Basque speakers are driven by default post-nominal suffix analysis to bootstrap syntactic cues in their mother tongue (Molnar et al., 2014), and they apply the same strategy in the L2.

This hypothesis implies two fundamental issues. First, it provides a step further in the research on multilingual prediction that goes beyond recent paradigms that have worked on the presence or absence of L2 prediction depending on the speakers’ proficiency. These results show that the native language experience strongly influences the way prediction is carried out, in other words, the environmental language regularities available during early childhood shape reliance on different levels of linguistic representations for prediction.

This leads to the second point: the experiment provides electrophysiological evidence that language prediction is not a “representation encapsulated” phenomenon (dealing only with high-level linguistic representations) but it flexibly takes advantage of the linguistic representations made available by different cognitive processes across the form-to-meaning (perception-to-abstraction) hierarchy. Predictive coding approaches (Bastos et al., 2012; Friston, 2005; Rao & Ballard, 1999) assume that human interaction with the environment is largely based on internal knowledge-based expectations (*priors*) that, through a top-down process, hierarchically percolate down from memory-based abstract internal representations to sensory regions, thus shaping human perception. So far, experimental evidence of such prediction hierarchy in the language domain has been missing and our recent data (Molinaro et al., 2016; under review) provides initial evidence in this direction.

### The present study

The aim of the present study is to test the above hypothesis concerning hierarchical predictions during language processing (Molinaro et al., 2016; under review). In particular, we sought for additional evidence that prediction processing is actually hierarchical, by testing the format of the representation on which prediction is based.

We analysed two similar groups of highly proficient bilinguals: participants were Spanish (L1) – Basque (L2) speakers and Basque (L1) – Spanish (L2) speakers, with the L2 acquired at the age of three years old. In contrast with the previous experiments, we decided not to use sentences providing a constraining context to trigger lexical prediction. In a sentence context, the effects recorded on the article do not ensure that participants are actually predicting the following content word, but could reflect prediction of the determiner (see Luke & Christianson, 2016), as it plays a relevant sentence-level syntactic role. We thus utilized a word matching task (referred to as WMT in the following) and a picture matching task (PMT), and combined them with the EEG to have the necessary high-temporal resolution to detect anticipation effects.

In two different blocks of the same experimental session, participants read a noun in Spanish on a screen, or they saw the picture of a noun on the screen (e.g. *cuchillo* (knife) or the image of a knife). These stimuli (referred to as *predictors*) were followed by a voice saying a noun phrase (NP, a noun – the *predictee* – preceded by a determiner). Here the noun could be congruent or incongruent with the word that they read or the image they saw. The gender of the incongruent noun was the opposite of the gender of the presented stimuli; furthermore, nouns could be transparent or opaque. The determiner preceding the noun in the NP could be congruent, incongruent or neuter in relation to the stimuli, but it was always in agreement with its noun. Importantly, the determiner could also be gender-congruent with the predictee but it preceded a mismatching noun in some trials.

Participants had to indicate whether what they read or what they saw (the predictor) corresponded to what they heard (the predictee).

We recorded the ERPs time-locked to both the determiner and the noun that the participant heard.

The reasoning behind the experiment comes from straightforward psycholinguistics models.

In language processing, when users read a word, sensory information feedforwards activation to the sub-lexical (abstract orthographic) word-form features of that item (reliably after 200 ms post-stimulus onset), and then to the lexical/semantic information related to that word (after 350 ms, Grainger & Holcomb, 2009; for theoretical proposals see Grainger & Jacobs, 1996; McClelland & Rumelhart, 1981). In contrast, after picture presentation, the visual information directly activates the lexical/semantic information related to the image (after 200 ms post-stimulus), from which inflected word-forms can be derived (after 300 ms), similarly to what happens during language production (Indefrey & Levelt, 2004; Strijkers & Costa, 2016; for a parallel between prediction and production see Dell & Chang, 2014; Pickering & Garrod, 2013; Molinaro et al., 2016).

The rationale of the present study is to track the time course of the mismatching effect (if we observe any) time-locked to the determiner, depending on the predictor format (either a written word or a picture). In order to perform the matching task, participants pre-activate the word they expect to hear (the predictee), according to the item they perceived (the predictor). The gender values of the predictor will be consequently activated in time depending on the hierarchical level in which they are represented. In the case of WMT, we expect that the predictor (the written word) will sequentially activate the following representational levels, i.e., visual > sub-lexical > lexical. Importantly, access to lexical representation is mediated by sub-lexical processing. If the gender value is represented at the sub-lexical level, the mismatching gender information provided by a mismatching determiner will trigger an update of the prediction earlier in time compared to a lexical-level gender representation. The effect of mismatch between the expected determiner and the perceived determiner represents the electrophysiological correlate of prediction.

In contrast, in the PMT, the feature pre-activation process triggered by the predictor (the picture) would be visual > lexical > sub-lexical^1^; in other words, full lexical access occurs earlier since it is not mediated by sub-lexical units. If the gender value is represented at the lexical level of processing it will trigger an earlier prediction update compared to a scenario in which gender is strictly sub-lexically encoded. Still, since gender can be reliably represented through both a lexical and a form-based route (Gollan & Frost, 2001), there would be no special need of activating sub-lexical representations in the PMT since gender values would be already lexically available.

Crucially, our experiment did not only manipulate gender but also transparency. Let us consider the opaque words first. In order to predict the gender of opaque nouns participants need to rely on the lexical information, as there are no formal cues to give them clues about the gender. Therefore, in both groups of speakers we expect the unexpected determiner to elicit the same pre-activation timing as found in the previous experiments (the effect should be earlier in the PMT than in the WMT since the lexical information is earlier available in the former). The timing of opaque words represents our reference point for lexical prediction.

On the other hand, transparent words should display a different pattern in the two groups. In line with the previous experiment, we assume that Spanish (L1) speakers rely on lexical information for the pre-activation of gender, so we expect them to show a lexical effect (with similar timing to the opaque items) after they hear an unexpected article, both in the WMT and in the PMT.

In contrast, we predict Basque (L1) natives performing the WMT to present an earlier effect on the unexpected determiner (compared to Spanish natives), as they rely on sub-lexical information for the pre-activation of gender. When they see an image in the PMT, the lexical information will be accessed before the sub-lexical one. Thus, the gender would be pre-activated through the lexical route, but it would also be followed by an additional sub-lexical analysis. As a consequence, we expect Basque natives to show a delayed effect in comparison to Spanish natives.

If our expectations are correct, we will have further evidence that there are hierarchical levels of representation in language prediction; that prediction mechanisms can navigate this hierarchy quickly adjusting to the input; and that the language acquired first has an impact on language anticipation processes, even in early bilinguals with very high proficiency. This would add important information to language prediction research, and provide new insights into the mechanisms operating during monolingual and bilingual language processing.

## Material and methods

### Participants

Forty-two early bilinguals participated in the experiment. They were divided in two groups. The first group was formed by twenty-one native speakers of Spanish who were first exposed to Basque after the age of three (11 females; age range 19–29, mean: 23.38, SD: 3.24: Age of acquisition of Basque: 3.61 y.o., SD: 1.46).

Twenty-one native speakers of Basque (13 females; age range 18–33, mean: 25.66, SD: 5.45: Age of acquisition of Spanish: 4.23 y.o., SD: 1.33) who started to learn Spanish after the age of three formed the second group. All participants were right handed, their vision was normal or corrected to normal, and they had no history of neurological disorder. Before taking part in the experiment, participants signed an informed consent. They received a payment of 10 € per hour for their participation. The study was approved by the BCBL ethics committee.

In order to participate in the study, all the participants had to go through some language proficiency tests in both Spanish and Basque (results in Table 1). First, participants had to self-rate their language comprehension (on a scale from 1 to 10, where 10 was a native-like level; the result was averaged for speech comprehension, speech production, reading and writing). Basque speakers rated themselves very high in both Basque and Spanish, while Spanish speakers claimed they were better in Spanish than Basque.

**Table 1:** General proficiency assessment of the participants in the two groups

| **Measure** | **Spanish natives (N=21)** | **Basque natives (N=21)** |
| --- | --- | --- |
| Self-evaluation (0-10) |  |  |
| Spanish | 9.37 (0.29) | 9.39 (0.24) |
| Basque | 8.02 (0.49) | 9.04 (0.16) |
| LexTALE |  |  |
| Spanish (0-60) | 52.76 (5.21) | 54.09 (4.13) |
| Basque (0-50) | 34.38 (5.88) | 46.04 (2.67) |
| Picture naming (0-65) |  |  |
| Spanish | 64.47 (0.92) | 63.38 (1.62) |
| Basque | 50.09 (9.63) | 64.19 (1.47) |
| Interview (0-5) |  |  |
| Spanish | 5 (0) | 5 (0) |
| Basque | 4.14(0.47) | 4.95(1.33) |

Participants also went through a lexical decision task called LexTALE (Izura et al., 2014; Lemhofer & Broersma, 2012) to test their vocabulary knowledge. In Spanish, both groups showed the same very high score, but in Basque, Spanish speakers had lower scores than Basque natives.

Then participants had to name a sequence of pictures which increased in difficulty, in both languages. Again, both Spanish and Basque participants had the same native-range score in Spanish, but Basque speakers were better than Spanish speakers in Basque. Finally, all the participants were interviewed by balanced bilingual linguists who rated them on a scale from 0 to 5: in both languages no participants had a score below 4. All the participants studied English at school, and claimed it was their third language.

We applied the same measures to test subjects’ English proficiency: there was no difference between groups, scores were good, but still much lower than Spanish and Basque (self-evaluation: 5.2, SD: 3.22; LexTALE, score 0−40: 22.43, SD: 5.15; picture naming: 44.07, SD: 7.22; interview: 3.88, SD: 1.44). Since participants were far more proficient in their two other languages than in English, we assumed that this third language could not influence the present design.

### Experimental design and materials

A list of 120 Spanish nouns was selected, where 60 nouns were transparent and 60 nouns were opaque (30 masculine and 30 feminine nouns per group).

The transparent masculine nouns ended with “*-o*”, which is the typical Spanish ending for masculine, (e.g. *cuchillo*, “knife [masc]”), while the feminine nouns had the feminine ending “*-a*” (e.g. *silla*, “chair [fem]”). Irregular nouns were excluded. Opaque nouns showed endings that were not informative of the grammatical gender (i.e., “*-e*”, “*-n*”, “*-l*”, “*-s*”, “*-j*”, “*-r*”, “*-d*”, “*-z*“).

The mean number of letters for transparent and opaque nouns was identical (mean: 5.76 letters, SD 1.51; range: 4–9 letters). In addition, transparent and opaque nouns did not differ for measures of concreteness (transparent nouns mean: 5.86, SD 0.58; opaque nouns mean: 5.67, SD 0.63); imageability (transparent nouns mean: 6.11, SD 0.73; opaque nouns mean: 6.09, SD 0.75) and familiarity (transparent nouns mean: 6.16, SD 0.42; opaque nouns mean: 6.12.67, SD 0.53) (EsPal, Duchon et al., 2013).

Half of the nouns referred to artifacts, and half to natural objects.

The words selected were also used to create the stimuli for the picture matching task. We found images that visually represented the nouns above. All the pictures were highly recognizable colour photographs (.png extension, white background, 2000x2000 pixels) obtained from online image collections. In order to be sure that a picture could only be related to one possible noun, we ran a naming test. Spanish-Basque bilinguals (N=20) who did not take part in the experiment saw 240 images, and named them with the first noun that came up to their minds. We only chose the images whose name was univocally expressed by all the 20 participants, and we came up with a final 120 pictures that could only represent our original list of nouns. Both the words and the images were followed by an auditory noun phrase formed by a determiner followed by a noun. The determiner could be a definite article (*el* “the [masc]”, *la* “the [fem]”), or a possessive adjective (*mi* “my”, *su* “his, her”) that was gender unmarked, hence neuter. The noun could either match or mismatch with the previously presented stimulus, but always matched with its own determiner. The experiment had 5 conditions. There were 2 main conditions (that represented 53% of the trials). In the first one, both the determiner and the noun matched with the stimuli (written or visual); in the second, both the determiner and the noun mismatched with the stimuli. The gender value of the determined was balanced in the two conditions. In addition, we added a condition (26% of the trials) in which the determiner was neuter, and the noun matched with the stimuli. This last condition was introduced to reduce strategical prediction effects in our experimental design time-locked to the determiner. However, since the sound envelope of the neuter determiner was different compared to the experimental ones, we reasoned that the relative evoked response could not be directly compared with the experimental ones. Furthermore, there were 2 catch trials conditions (20% of the trials): in the first, a neuter determiner was followed by noun that mismatched with the previous stimulus; in the second, the determiner matched with the stimulus, but the noun did not.

For each word/image of the original list, 5 possible noun phrases were recorded, corresponding to each of the 5 conditions (e.g. *BOTELLA* “bottle [fem]”: *la botella* “the [fem] bottle”; *mi botella* “my bottle”; *el mando* “the [masc] remote control”; *su mando* “his/her remote control”; *la corona* “the [fem] crown”).

Sound strings were recorded by a Spanish female speaker. Between the determiner and the noun there was a silence gap of about one second (range 1- 1.3s, the exact timing was measured for each item). All the items were checked for amplitude (recording, cuts, measures and standardization was done using Praat (Boersma & Weenink, 2007).

In order to assess how many milliseconds a listener needs to distinguish between the determiner *el* “the [masc]”, and the determiner *la* “the [fem]”, we ran a discrimination test with 10 participants who did not take part in the experiment. We took five NPs whose determiner was *el*, and five whose determiner was *la*. The audio files corresponding to the determiners were cut in order to create smaller time windows. Participants listened to audio fragments containing the first 60 ms; 70 ms; 80 ms or 90 ms of the determiner of each NP for a total of 40 trials, and they had to indicate (by spelling it aloud to an experimenter) whether they thought it was an *el* or a *la*. For the 60 ms bits the inaccuracy percentage was 54% for the determiner *el*, and 64% for the determiner *la*. Participants listening to audio files lasting 70 ms had an inaccuracy of 16% for *el*, and 28% for *la*. When audio fragments lasted 80 ms participants recognized all trials with the determiner *el*, but they got wrong 3% of the *la* determiners. There was 100% of accuracy when the audio files lasted 90 ms for both determiners. Therefore 90 ms is defined as the uniqueness point (Marslen-Wilson, 1987) of the determiner gender value, and will be later subtracted from the emerging ERP component timing. For both the WMT, and the PMT, 3 lists were created.

Each list had 240 visual stimuli, each of the 3 main conditions had 64 items, and the catch trial conditions had 24 items each. In each list, visual stimuli were repeated twice, but never in the same condition. Participants were never given the same list for both WMT and PMT, furthermore, the task sequence was alternated among participants, so that half of them first went through the WMT and then the PMT, and the other half did the opposite.

Within each list the words were balanced (all *p* > 0.2; based on EsPal, Duchon et al., 2013) for grammatical gender, word frequency (log-values: List1: 1.21, SD: 0.55; List 2: 1.26, SD: 0.52; List 3: 1.25, SD: 0.56), and number of letters (List1: 5.79, SD: 1.59; List 2: 5.79, SD: 1.46; List 3: 5.68, SD: 1.51). Finally, all the images were balanced among the lists, so that they were all equally distributed in all the conditions. No differences emerged between lists.

### Procedure

The EEG experiment was run in a soundproof electrically shielded chamber with a dim light. Participants were seated in a chair, about sixty centimeters in front of a computer screen. Stimuli were delivered with the PsychoPy software (Peirce, 2007).

In the WMT, participants read words displayed in black letters on a white background. In the PMT, subjects saw images in the center of the screen. After a fixation cross (lasting 500 ms), the words or the images appeared on the screen for 350 ms. After the visual stimuli disappeared, they were immediately followed by auditory stimuli played by two speakers. The stimulus onset asynchrony (SOA) between visual and auditory stimulus was selected so that the onset of the auditory stimulus was time-locked to the on-going lexical processing of the visual word in the WMT and to sub-lexical processing in the PMT. In addition, studies employing cross-modal priming have shown that the effects tend to be more robust when the SOA is larger than 200 ms (Holcomb & Anderson, 1993). After participants heard the NP, they had to indicate whether the word they read or the image they saw matched with the noun they heard. The question appeared in the center of the screen as soon as the sound finished, and the subject could answer using the relative buttons on the keyboard: the response hand was counterbalanced across participants and list. Numbers of correct responses were recorded, and RTs were calculated in milliseconds from the appearance of the question to the participant’s key press. All trials were presented in a different random order for each participant. WMT and PMT were presented in two parts of the same experimental session, with a small break between the two.

A brief practice session included five words in one session, and five images in the other, followed by the relative auditory stimuli and the yes-no questions. Participants were asked to stay still and to try to reduce blinking and eyes movement to minimum, especially during the auditory presentation. Overall, the experiment lasted one hour on average.

### Electrophysiological recording and data analysis

EEG was recorded from 27 electrodes placed in an elastic cap (Easycap, www.easycap.de): Fp1, Fp2, F7, F8, F3, F4, FC5, FC6, FC1, FC2, T7, T8, C3, C4, CP5, CP6, CP1, CP2, P3, P4, P7, P8, O1, O2, Fz, Cz, Pz. All sites were online referenced to the left mastoid (A1). Additional external electrodes were placed on mastoids (A1, A2) and around the eyes (VEOL, VEOR, HEOL, HEOR) in order to detect blinks and eye movements. A forehead electrode served as the ground. Data were amplified (Brain Amp DC) with a bandwidth of 0.01−100 Hz, at a sampling rate of 250 Hz. The impedance of the scalp electrodes was kept below 5 kΩ, while the eye electrodes impedance was below 10 kΩ. Collected recordings were off-line re-referenced to the average activity of the two mastoids. Artifacts exceeding 100 μV in amplitude were rejected. Raw data were visually inspected and artifacts such as muscular activity and ocular artifacts were marked for subsequent rejection. On average, 7.3% of epochs were excluded as considered artifacts. There was no difference between conditions and groups in terms of artifact rejection.

For the analysis of the determiner, epochs of 1600 ms (from –600 ms to 1000 ms) were obtained, considering a –600 ms pre-stimulus baseline. For each condition, the average ERP waveforms were computed time-locked to the onset of the determiner. Epochs were averaged independently for each condition and subject. In order to obtain a detailed exploration of the exact time course of the effects on the determiner, and define their evolution in time, we performed pairwise comparisons of the ERP waveforms (match vs. mismatch) with point-by-point (one point every 4 ms) t-test for each electrode. We ran separate comparisons for each task, each gender type and each language group. To protect this analysis from false positives we employed the Guthrie and Buchwald (1991) correction that filters out effects which last less than 50 ms (12 consecutive time points) in less than three sensors. Importantly, this approach does not constrain the selection of the time interval of interest. However, we validated the relevant point-by-point effects that could reflect an interaction between the main factors of interest (match by transparency) within each Experiment (Word/Picture matching task) and within each group (Basque/Spanish natives) with further statistics. We selected the 100 ms-long time interval of interest and entered the average ERP activity – across the electrodes in which a significant effect emerged – in a three-way ANOVA with Match (two levels: match vs. mismatch), Transparency (two levels: transparent vs. opaque) and Electrode (variable number of levels depending on the electrodes of interest). P-values were Greenhouse-Geisser corrected.

For the analysis of the noun, we computed 3100 ms epochs (from –1600 ms to 1500 ms), applying the same pre-stimulus baseline used for the determiner, so that we could have all the NP electrophysiological time course, but the waveforms were time-locked at the exact onset of the noun for each condition. Here, the statistics time-locked to the noun focused on the time-interval classically related to lexical integration processing (300–700 ms, namely the N400).

## Results

### Behavioural

Inaccuracy (mean percentage of incorrect responses) and reaction times (RT) from accurate trials were analyzed.

In WMT, Spanish natives had a general average inaccuracy of 4.50 % (6.54% for the opaque words, and 2.46% for the transparent words). In Basque natives the average inaccuracy was 3.29 %(5.03% for opaque words, and 1.54% for the transparent words).

For the PMT, there was 3.11% of inaccuracy in Spanish natives (4.20% for the opaque words, and 2.02% for the transparent words). Basque natives gave 6.98% of inaccurate answers (8.01% for the opaque, and 5.95 for the transparent).

In the Spanish group, the mean reaction time for the WMT was 490 ms (SD: 0.37), while for the Basque groups it was 500 ms (SD: 0.30). In the PMT, Spanish natives has 490 ms (SD: 0.50) reaction time, and for the Basque natives it was 470 ms (SD: 0.24).

To analyze the reaction time values of the accurate trials for the WMT and the PMT we used two mixed-design ANOVAs, with Transparency and Condition as within factors, and with Group as between factor. No significant differences emerged from the analysis: there were neither main effects, nor interactions among variables (Table 2.)

**Table 2:** ANOVA results of WMT and PMT reaction time values

| Measure | WMT | PMT |
| --- | --- | --- |
| Condition | $F(4, 52) = 1.36, p = .251, \eta_p^2 = .43$ | $F(4, 52) = .62, p = .649, \eta_p^2 = .16$ |
| Transparency | $F(1, 38) = .76, p = .389, \eta_p^2 = 0.20$ | $F(1, 38) = .29, p = .596, \eta_p^2 < .01$ |
| Group | $F(1, 38) = 1.36, p = .758, \eta_p^2 = .03$ | $F(1, 38) = .33, p = .567, \eta_p^2 < .01$ |
| Condition*Group | $F(4, 52) = 1.32, p = .265, \eta_p^2 = .37$ | $F(4, 52) = .79, p = .531, \eta_p^2 = .20$ |
| Transparency*Group | $F(1, 38) = 2.16, p = .150, \eta_p^2 = .54$ | $F(1, 38) = 89, p = .353, \eta_p^2 = .23$ |
| Condition*Transparency | $F(4, 52) = 1.34, p = .260, \eta_p^2 = .24$ | $F(4, 52) = .54, p = .706, \eta_p^2 = .14$ |
| Condition*Transparency*Group | $F(4, 52) = .31, p = .869, \eta_p^2 < .01$ | $F(4, 52) = .60, p = .662, \eta_p^2 = .01$ |

### ERP data

#### Determiner

In order to better highlight the prediction effects time-locked to the determiner presentation, we report data in the time window until 1 sec. In the Figures below (1–4) we report the ERPs for the electrodes in the left hemisphere scalp region in which significant effects emerged; the point-by-point analyses showing activation timing across all the electrodes; and the scalp topography relative to the 250–350 ms and 400–500 ms time windows.

**Figure 1:**
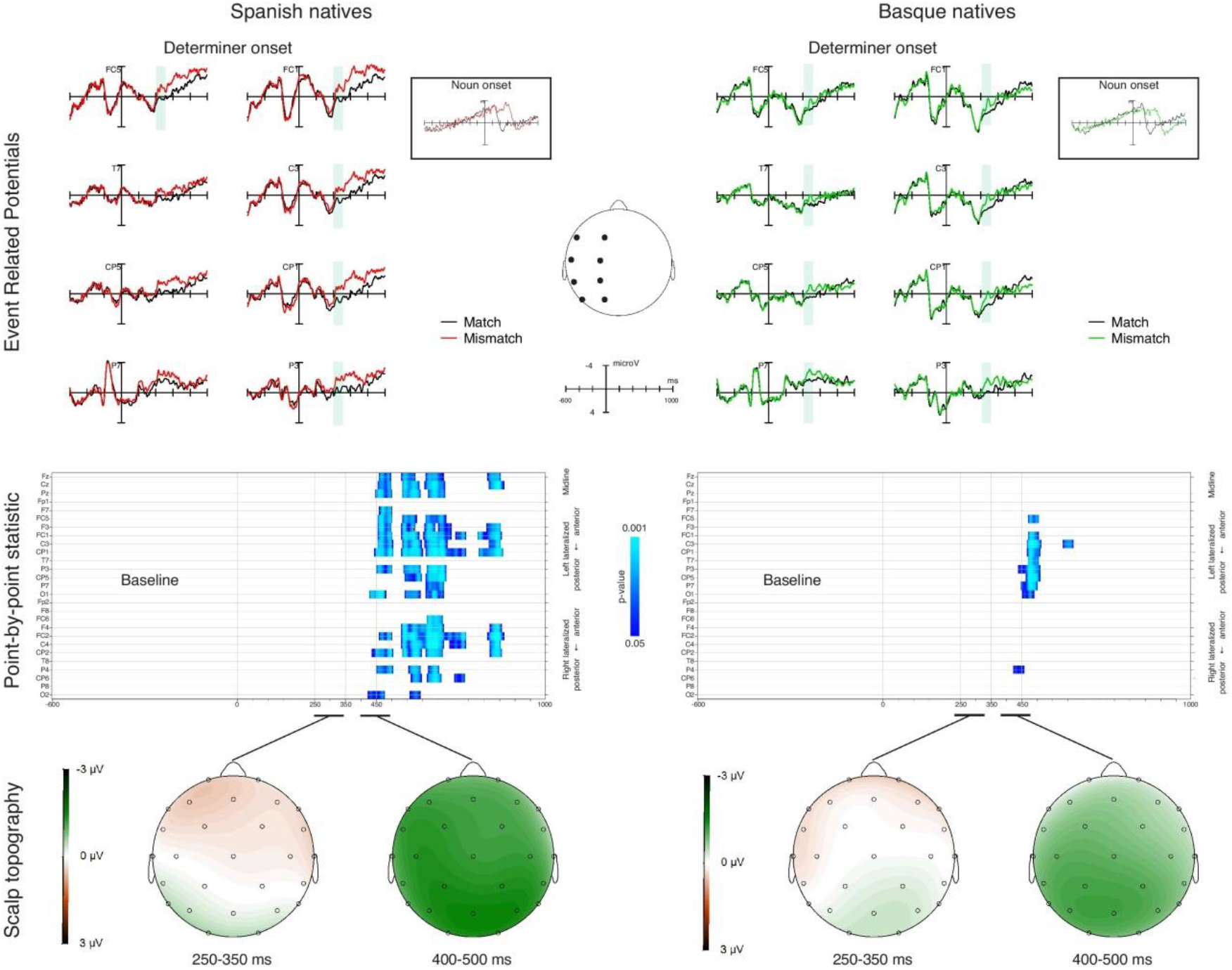
ERPs for the **WMT** relative to the match and mismatch condition (Prediction effect) time-locked to the presentation of the determiner preceding the predicted **opaque** noun (plotted in –600 to 1000 for better display of the ERP effects). We separated the two groups of participants and plotted all the electrodes. Shadowed differences are the statistically significant ones relative to the 250–350 ms time window. The rectangles on the right side of the graphs present the grand average ERPs relative to the match and mismatch condition (Integration effect) time-locked to the presentation of the **opaque** noun (plotted in –1600 ms to 1500 ms), for all the electrodes. The point-by-point plot analysis of variance for each electrode (Guthrie and Buchwald, 1991, corrected) is presented below. We report the main effect of the Prediction factor: vertical blue lines indicate the interval in which the effect emerged. In the lower panels we present the topographical distribution of the difference effect (mismatch minus match) in the two time intervals of interest (250–350 ms; 400–500 ms).

**Figure 2:**
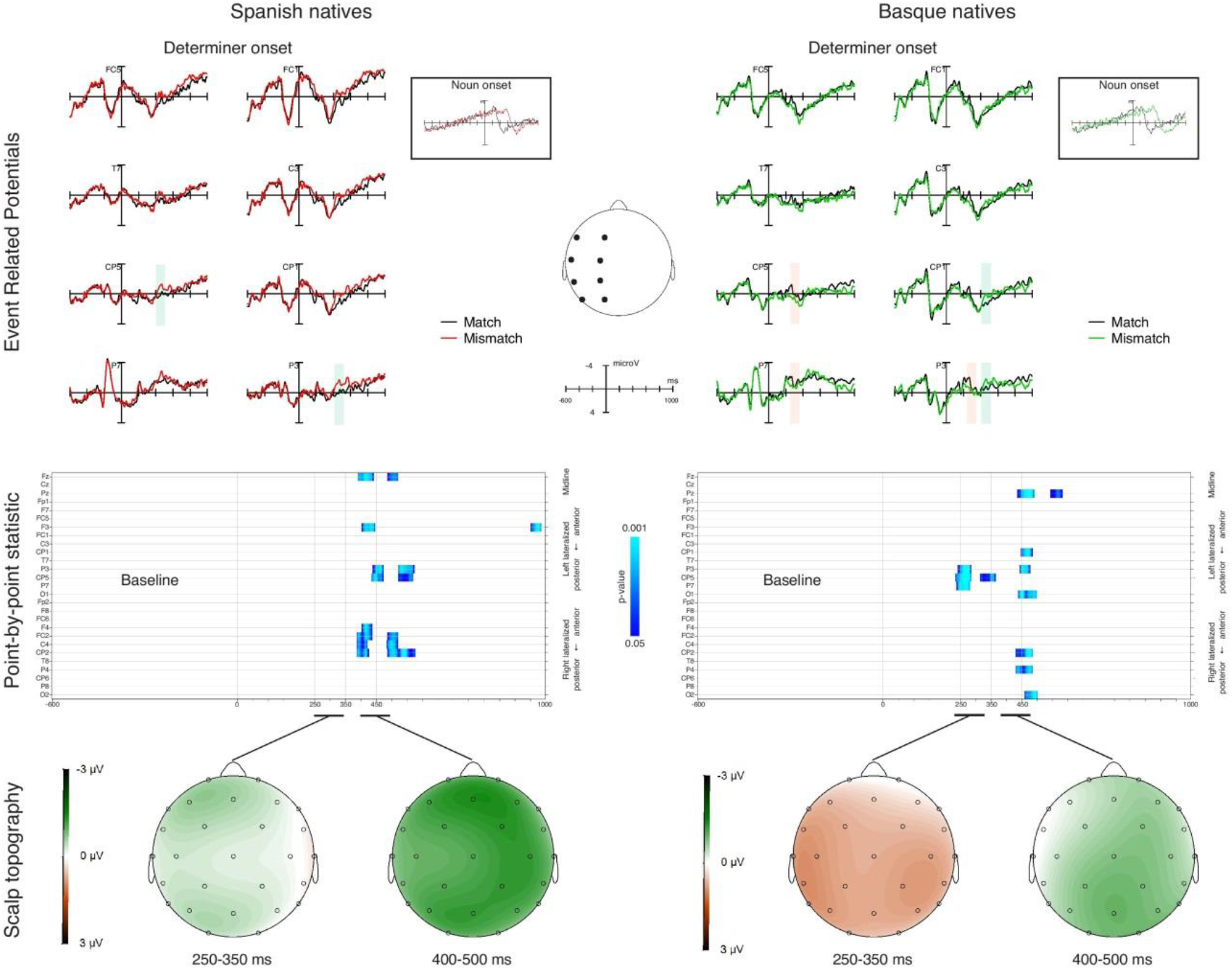
ERPs for the **WMT** relative to the match and mismatch condition (Prediction effect) time-locked to the presentation of the determiner preceding the predicted **transparent** noun (plotted in –600 to 1000 for better display of the ERP effects). We separated the two groups of participants and plotted all the electrodes. Shadowed differences are the statistically significant ones relative to the 250–350 ms time window. The rectangles on the right side of the graphs present the grand average ERPs relative to the match and mismatch condition (Integration effect) time-locked to the presentation of the **transparent** noun (plotted in –1600 ms to 1500 ms), for all the electrodes. The point-by-point plot analysis of variance for each electrode (Guthrie and Buchwald, 1991, corrected) is presented below. We report the main effect of the Prediction factor: vertical blue lines indicate the interval in which the effect emerged. In the lower panels we present the topographical distribution of the difference effect (mismatch minus match) in the two time intervals of interest (250–350 ms; 400–500 ms).

**Figure 3:**
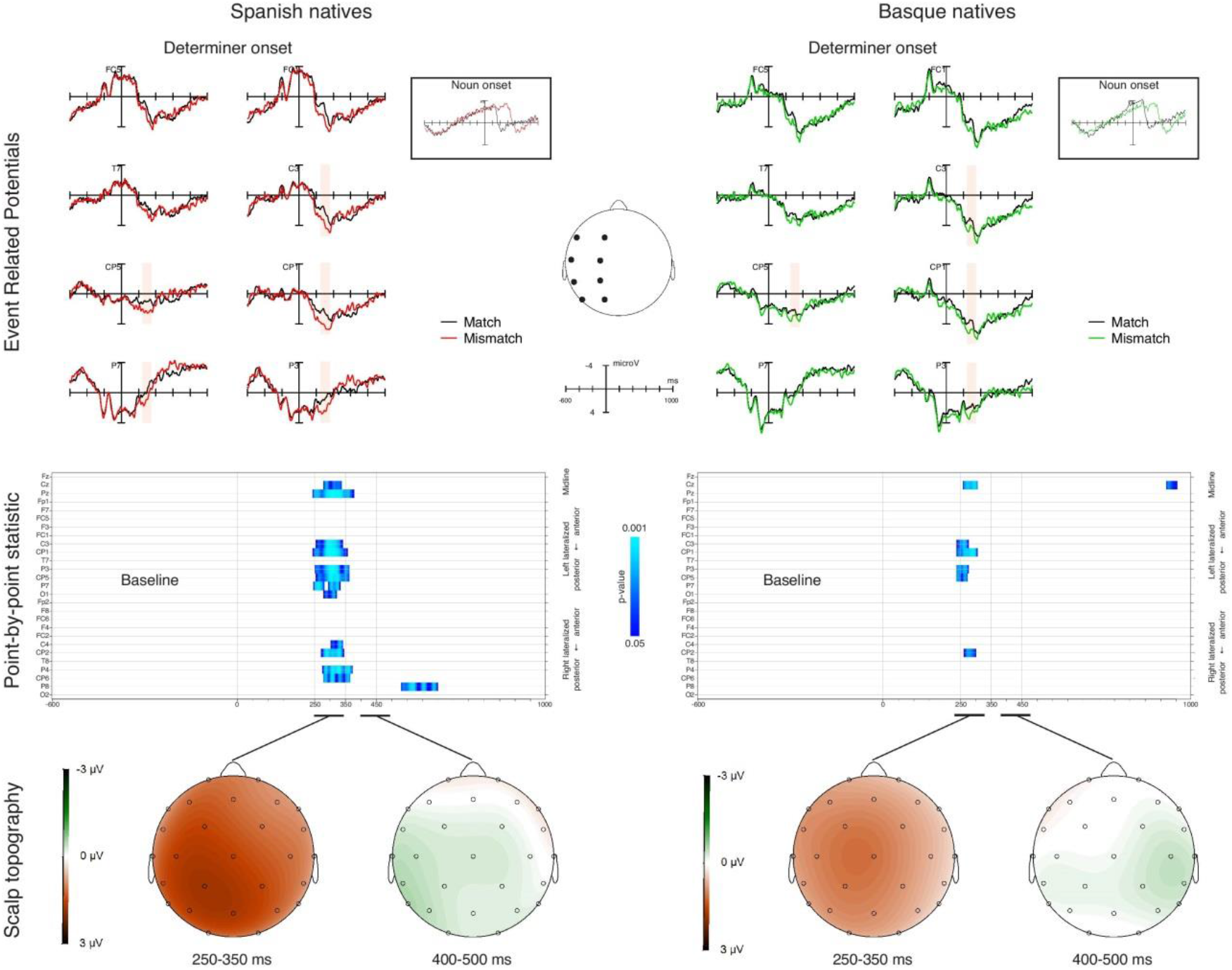
ERPs for the **PMT** relative to the match and mismatch condition (Prediction effect) time-locked to the presentation of the determiner preceding the predicted **opaque** noun (plotted in –600 to 1000 for better display of the ERP effects). We separated the two groups of participants and plotted all the electrodes. Shadowed differences are the statistically significant ones relative to the 250–350 ms time window. The rectangles on the right side of the graphs present the grand average ERPs relative to the match and mismatch condition (Integration effect) time-locked to the presentation of the **opaque** noun (plotted in –1600 ms to 1500 ms), for all the electrodes. The point-by-point plot analysis of variance for each electrode (Guthrie and Buchwald, 1991, corrected) is presented below. We report the main effect of the Prediction factor: vertical blue lines indicate the interval in which the effect emerged. In the lower panels we present the topographical distribution of the difference effect (mismatch minus match) in the two time intervals of interest (250–350 ms; 400–500 ms)

**Figure 4:**
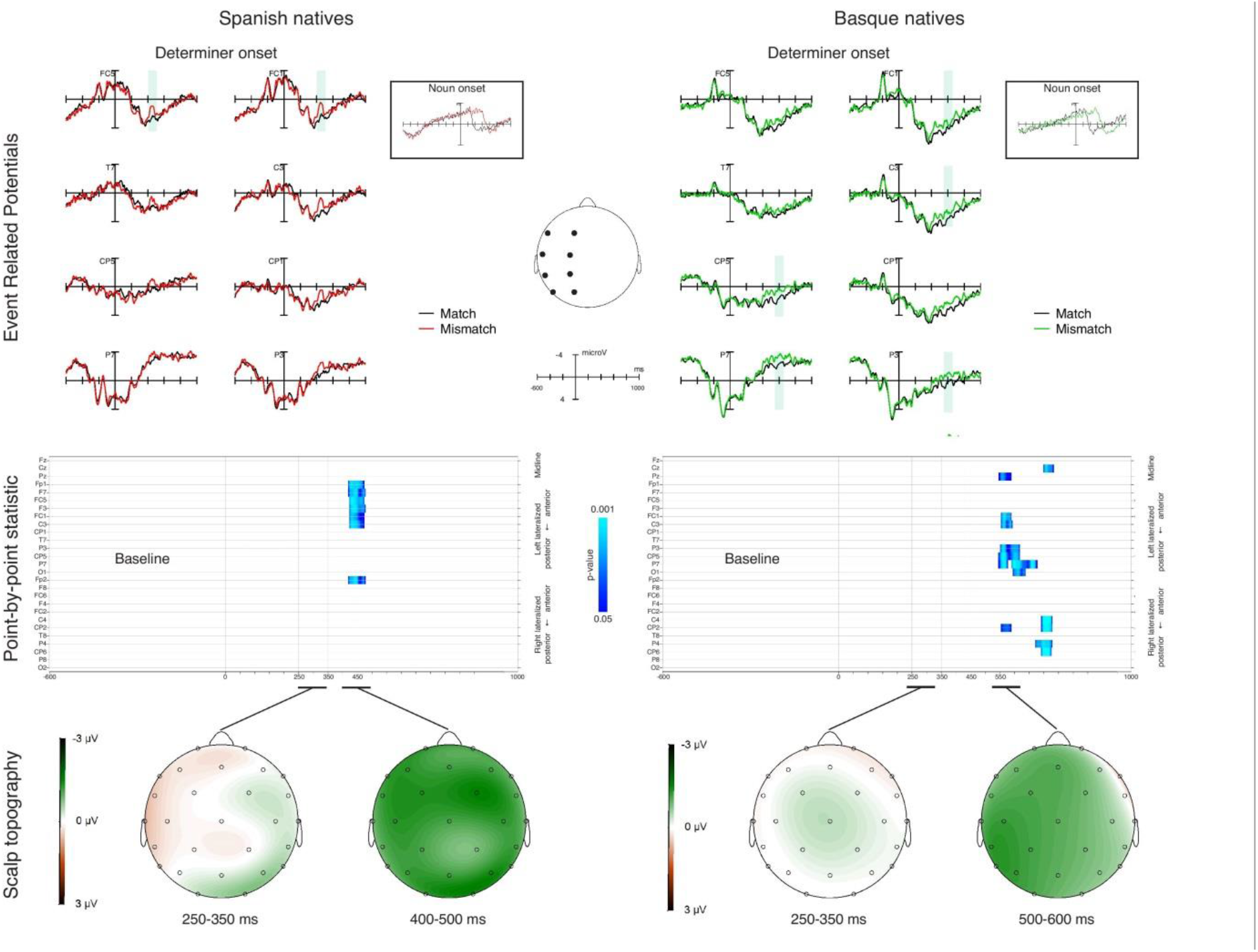
ERPs for the **PMT** relative to the match and mismatch condition (Prediction effect) time-locked to the presentation of the determiner preceding the predicted **transparent** noun (plotted in –600 to 1000 for better display of the ERP effects). We separated the two groups of participants and plotted all the electrodes. Shadowed differences are the statistically significant ones relative to the 250–350 ms time window. The rectangles on the right side of the graphs present the grand average ERPs relative to the match and mismatch condition (Integration effect) time-locked to the presentation of the **transparent** noun (plotted in –1600 ms to 1500 ms), for all the electrodes. The point-by-point plot analysis of variance for each electrode (Guthrie and Buchwald, 1991, corrected) is presented below. We report the main effect of the Prediction factor: vertical blue lines indicate the interval in which the effect emerged. In the lower panels we present the topographical distribution of the difference effect (mismatch minus match) in the two time intervals of interest (250–350 ms; 400–500 ms for the Spanish natives and 500–600 ms for Basque natives).

In the WMT, for the Spanish group, determiners preceding the opaque nouns display a more negative effect for Mismatch items, likely reflecting a N400 modulation: this long-lasting effect starts at about 420 ms (330 ms post-uniqueness point), with a widespread distribution in all the electrodes. Basque natives showed a similar short-living negative effect starting 430 ms (340 ms post-uniqueness point).

The WMT relative to the determiners preceding transparent nouns showed different results. Spanish natives display a strong effect of Prediction in the time interval starting around 380 ms (290 ms post-uniqueness point): the effect is more negative for Mismatch compared to Match condition and it could be interpreted as an N400 evident in frontal and parietal electrodes.

Crucially, in the same condition Basque natives have an earlier effect in the time window from 240 ms to 350 ms (150 ms post-uniqueness point). The effect is most positive for the Mismatch condition and particularly evident on the left parietal electrodes, and it can be interpreted as a P200 effect.

The early effect is followed by an N400 effect starting at about 430 ms (340 ms post-uniqueness point) with a wider scalp distribution.

The three-way ANOVA run to check for robustness of Prediction effect in the 250–350 ms time window (for the electrodes P7, P3, and CP5) in both groups showed an interaction between Match (congruency between stimulus and determiner) and Transparency only for Basque natives [Spanish natives: F(1,20)=0.63, p>0.4, ges=0.002; Basque natives: F(1,20)=5.01, p<0.05, ges=0.006]; importantly, it also showed an interaction between Match and Group [F(1,40)=3.89 p<0.05 ges=0.012].

The results of the PMT display a different pattern. For determiners preceding opaque nouns, in both Spanish and Basque natives there is a robust effect in an early time window, from 250 (160 ms post-uniqueness point) to 350 ms. The effect is more positive for mismatching items compared to matching items. We assume this to be an early lexical effect.

The effect on transparent nouns was different. In Spanish natives, determiners preceding transparent nouns elicit an increased negative effect for mismatching items at about 420 ms (330 ms post-uniqueness point). The effect is frontally distributed. A more left-posteriorly distributed negative effect for mismatching items is also present in Basque natives: it starts later, at 550 ms (460 ms post-uniqueness point). For the 250–350 ms time window, in the Spanish group, a three-way ANOVA showed interaction between Match and Transparency in the electrodes of interest [F (1,20) = 10.51, p < 0.01, ges = 0.011]. The Basque natives displayed the same interaction [F (1,20) = 4.65, p < 0.05, ges = 0.006]. In the later time intervals, we ran two separate ANOVAs considering Group and Match as separate factors. In the 550–650 ms time interval we observed a significant interaction between the two factors when considering the electrodes showing the significant difference for transparent items in the Basque group [F (1,40) = 3.60, p < 0.05, ges = 0.005]. However, no interaction was observed in the 400–500 ms interval (in contrast to the electrodes showing the significant difference for transparent items in the Spanish group) [F (1,40) = 1.66, p > 0.1, ges = 0.001]. We can thus conclude that Spanish natives showed a statistically reliable prediction effect in the 400–500 ms that was not strong enough to be reliable for Basques. In the later 550–650 ms time interval, a reliable prediction effect was observed in the Basque native group but not in the Spanish native group.

#### Noun

ERP results on the onset of the noun are clear-cut and straightforward. Given the long duration of the effect we found, the time window taken into consideration for the analysis is longer ending at 1.5 sec.

In both groups, both WMT and PMT produced a strong mismatch effect starting at about 300 ms and ending at about 800 ms. No significant differences were found between transparent nouns and opaque nouns. We assume this result to reflect a semantic integration effect.

We ran a three-way ANOVA with Match, Hemisphere and Longitude as factors, for the 300–700 ms time window and for all the electrodes. The analysis confirmed the results: the main effect of Match and the interactions between Match and Hemisphere and Match and Longitude are all significant (Table 3).

**Table 3:**
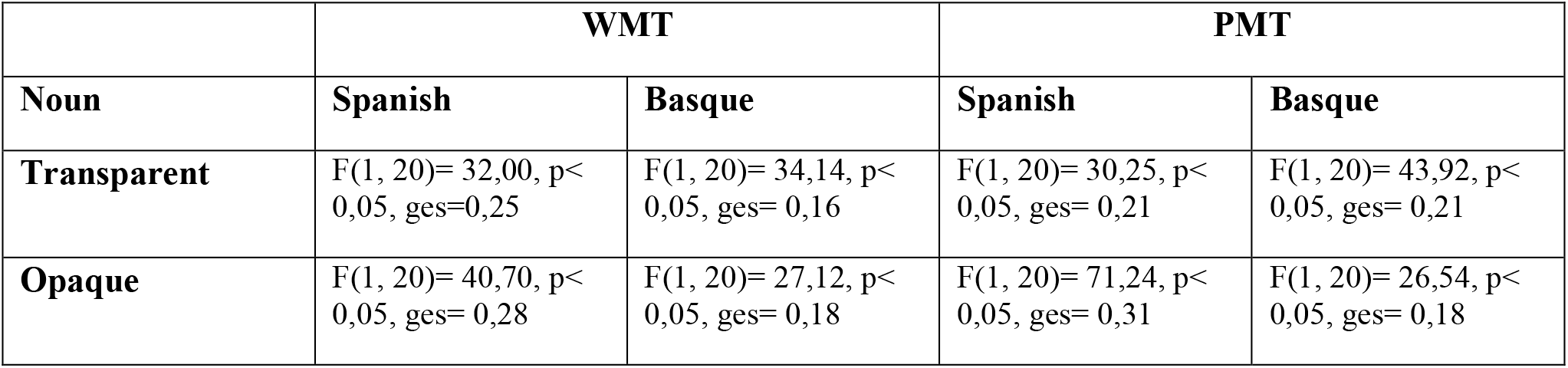
ANOVA main effects of Match on the noun (300–700 ms)

## Discussion

The goal of the present study was to provide evidence for the existence of hierarchical levels of representation in language prediction.

For this purpose, we used two groups of highly proficient bilinguals going through a WMT and a PMT. Subjects read a word, or saw a picture on a screen, followed by an auditory NP where the noun could be congruent or not with the preceding visual stimuli, and the determiner agreed with the noun. Also, the noun could be transparent or opaque.

The ERP effects recorded on the determiner preceding the noun (expected vs. unexpected determiner) revealed the time course of the processes operating inside linguistic prediction.

In the WMT, for the unexpected article preceding opaque words we found the same pre-activation timing for both Spanish and Basque natives. The effect started at ~430 ms (340 ms post-uniqueness point) in both groups, and it lasted longer in the Spanish group. This effect can be interpreted as an N400. The result is in line with our expectations: participants have to rely on their lexical knowledge (as reflected by the ERP lexical effect) for the gender prediction of opaque words, which do not provide formal cues about their gender.

The effects on determiners preceding unexpected transparent words showed different results. Spanish native ERPs revealed a negativity at ~390 ms (300 ms post-uniqueness point) roughly similar to the opaque words. This lexical effect was observed in Basque natives too, but crucially, they also displayed an early prediction effect at ~240 ms (150 ms post-uniqueness point). This confirmed our expectations: we had assumed that Basque natives rely more on sub-lexical, word-form related information for gender prediction of transparent words and we were expecting this prediction process to be reflected by the modulation of an early ERP component related to attentional perception, like the ~240 ms positivity found here. On the other hand, in Spanish natives predicting transparent words, we were expecting to see a lexical ERP effect (similar to the opaque words) as they would rely more on the lexical information for gender prediction, irrespective of the transparency of the target noun.

In the PMT, we assumed the lexical information to be available earlier compared to the WMT, as lexical access triggered by the picture would not be mediated by the sub-lexical units of the word, and would be direct. For the unexpected determiners preceding opaque words we found a similar effect at ~250 ms (160 ms post-uniqueness point) for both Spanish natives and Basque natives. We assume that this effect reflects a lexical/semantic process (if we assume the prediction/production parallelism, this “lexical timing” is typically reported in picture naming studies: Indefrey & Levelt, 2004; Strijkers & Costa, 2016). Here our expectations were confirmed, as the gender of opaque words can only be accessed and predicted through lexical information.

As predicted, the ERPs recorded on determiners preceding transparent nouns displayed an effect at ~430 ms (340 ms post-uniqueness point) in the Spanish group, and later effect (~550 ms, 460 ms post-uniqueness point) in the Basque group. Basque natives presumably performed additional sub-lexical analysis after the lexical one, resulting in later timing. Besides, in comparison to the determiners preceding opaque words, the effect happens later in time in both groups: a possible explanation for this dissimilarity will be given later in the discussion.

### The role of native experience

This experiment provides further evidence that lexical prediction does not involve predicting only the semantic traits of a word (Federmeier, 2007, Federmeier et al., 2002), but also its grammatical features. These features can be predicted even when they are not present in the bilinguals’ L1, in support of the idea that there are no separate domains of prediction, but that prediction is a unified mechanism looking for informative cues independently of the language processed.

Nonetheless, a central point emerging from the study is that language prediction is tuned to the characteristics of the L1 and consequently is extremely sensitive to the native experience. As the only difference between groups is the language learned first, we can assess that the prediction differences found between Basque and Spanish natives are due to the early L1exposure. During the first three years of their lives (in which they were only exposed to the L1) Basque speakers developed their prediction mechanisms following the regularities of Basque language. These mechanisms still influence Basque natives’ prediction processes even if they have been daily exposed to the statistical regularities of Spanish all the rest of their lives. On the other hand, speakers originally tuned to Spanish process transparent and opaque items equally during prediction, as a result of the more robust mapping of grammatical gender from lexical information in this language. This had already been found to be true not only in experiments with words in isolation (Caffarra et al., 2014), but especially in experiments investigating the effect of transparency during determiner-noun agreement (Caffarra & Barber, 2015). It is worthy of notice that the difference in prediction mechanisms between the two groups cannot be due to a reduced proficiency for Basque natives in Spanish, since in the general proficiency assessment the two groups were perfectly balanced in this language, and also the behavioral results in both tasks did not reveal any significant differences.

### Hierarchical levels of prediction

More importantly, this study provides stronger evidence for the existence of cognitive hierarchies in language prediction. Basque natives’ neurophysiological responses display two levels of language prediction in the WMT. At one level they rely on the sub-lexical information of the word that is going to come next, in order to predict its gender. At another stage, gender can be predicted according to the lexical information they have about that word (as stored in the mental lexicon). The two levels are differently represented in terms of neural timing, with the first operating earlier, at about 150 ms after gender disambiguation of the determiner, and the second operating later, in the N400 time window, as clearly emerged from the WMT. This result supports what we recently observed during sentence reading (Molinaro et al., 2016; under review), where both groups showed a lexical N400-like effect for lexical prediction during sentence reading, while the Basque natives showed an earlier P200-like effect only for the pre-activation of transparent words. Also, we have replicated this outcome using a completely different task, as we did not use highly constraining contexts in order to elicit prediction, as in the previous experiment, but we used a WMT where prediction was evoked by a single word on a screen. This is evidence that the kind of representation pre-activated in both Molinaro et al. (2016; under review) and in the WMT is strictly word-related. In addition, the prediction effects that we found are observable independently of the kind of experimental setting, supporting the idea that prediction-mechanisms are able to quickly and flexibly adjust to the different tasks a language user has to go through. Furthermore, in the present study, the predictees were not visual but were audio, providing critical evidence that prediction mechanisms are modality independent. The existence of hierarchies clearly emerges from the WMT, but what can the PMT tell us about them?

### An interface where hierarchical levels communicate

Given the characteristics of the PMT, in our opinion, a good way to approach and discuss the PMT results would be through psycholinguistic production models. According to these, the mental lexicon is conceived as a network in which concept, lemma (word stems) and lexeme (morphological units) are represented as spreading-activated, sequential, independent nodes (Roelofs, 1992; Levelt et al., 1998). It is feasible to assume that, in order to perform the PMT (and ultimately predict the word that is going to come with its relative gender), participants exposed to a picture activate conceptual/semantic information, which is directly followed by the activation of the lemma level representation; lastly they activate the information at the lexeme level.

In the PMT, unexpected determiners preceding opaque words elicited a lexical ERP effect at ~160 ms in both Spanish and Basque natives. On the other hand, unexpected determiners preceding transparent words produced a much later ERP effect, at about 330 ms for Spanish natives, and ~460 for Basque natives. The hierarchical spreading activation dynamics would be the same for both transparent and opaque words. Nonetheless, the two groups respond differently in the two cases, with the opaque words showing a much earlier activation compared to the transparent words. Why do the two groups display a difference in the prediction time course of transparent and opaque words? Also, why is there a difference in timing between groups only on determiners preceding transparent words?

One way to explain the different timing would be to suppose that the prediction of opaque words is based on the lemma representation participants have about those words, while for the prediction of transparent words participants rely more on the lexeme representation, so they predict the former earlier than the latter, as the former are activated before, in a cascade process.

A similar way to account for the difference found in the PMT between the prediction of transparent and opaque words would be to assume the existence of an interface between the lemma level and the lexeme level. The “160 ms effect” found on the determiner preceding the opaque words is a fully lexical effect. It could not be anything else because the gender of opaque words can only be extracted and predicted through the lexical route. In contrast, the later effect emerging at 340 ms in Spanish natives, and at 460 ms in Basque natives for articles preceding transparent words would be due to additional processing resources at a lower hierarchical level. We think that the gender of the transparent words cannot be predicted only with the lexical information, but needs (or simply is given) more support from the sub-lexical information level. This information is not directly available because of the nature of the PMT, and the resulting effect reflects an operation going on at the interface between lemma-related and lexeme, inflection-related information (visual > lexical > sub-lexical). This hypothesis is supported by the timing of the effect that, based on production models (Indefrey & Levelt, 2004; Strijkers & Costa, 2016), would reflect activation of sub-lexical phonological information.

Further support for the lemma-lexeme interface is given by the difference in timing that emerged on determiners preceding the transparent words, between Spanish and Basque natives. In the conflict at the interface level between the lemma features activated by the picture and the lexeme features accessed after it, Basque natives would rely more on sub-lexical analysis for the extraction and prediction of gender, while Spanish natives would depend on the support of the lexicon to perform the prediction. This process is reflected by an effect happening ~130 ms later than that of the Spanish natives.

The existence of an interface between lexical and sub-lexical knowledge that permits interaction between the two kinds of process might be partially visible also in the WMT. In this task we assumed the processing sequence to be visual > sub-lexical > lexical. As indicated above, the Spanish natives display a lexical effect at 290 ms on determiners preceding transparent words; this effect is also present for the determiner preceding opaque words in both groups, but it happens later, at 340 ms.

It is possible that the earlier lexical effect found in Spanish natives predicting transparent words in the WMT is due to the interface interaction between lexeme and lemma analysis. This interaction does not take place for the prediction of opaque words because their gender cannot be extracted through the sub-lexical information, and it does not happen in Basque natives predicting transparent words, as the first analysis they run is already fully sub-lexical. Unfortunately, the ANOVA looking for interactions between transparency, match and group did not show any significant value in this time window (290–340 ms), therefore there is no strong evidence supporting this idea.

Future studies can possibly shed light on the existence of the interface between lemma-related and lexeme-related information in language prediction.

### Results on the Noun

Finally, the ERPs recorded on the incongruent noun showed a strong late N400 effect. The effect was similar in both groups, in both tasks, in all the conditions. There was no significant difference and no interactions among the variables. This lexical effect is coherent with several previous studies on bilingual processes (Hahne, 2001; Hahne & Friederici 2001; Weber-Fox & Neville, 1996), and confirms that integration semantic processes are not influenced by the native language experience, especially when proficiency is very high. Also, this outcome is in line with the behavioural results, where no difference between groups was found.

Given the speakers’ capacity to mediate between native language experience, new language environment cues and task requirements, we assume prediction mechanisms to be extremely plastic and flexible.

## Conclusion

With the present set of data, we would like to emphasise the plasticity of language prediction mechanisms. This study not only shows that predictions can rapidly adapt to new contexts, selecting all the available cues in order to construct an internal representation of the new environment, but also that early experience models the way in which different cues are weighted to obtain optimal predictions. The whole process is extremely flexible, with neural timing constantly ready to adapt to the environmental demands and to mediate between native experience settings and new language feature requirements.

## Acknowledgments

This research was supported by the Spanish Ministry of Economy and Competitiveness (MINECO) [PSI2015–65694-P to N.M.]. The project was also partially supported by the award ‘‘Centro de Excelencia Severo Ochoa SEV-2015–0490”.

We wish to express our gratitude to the BCBL lab-staff and the research-assistants who helped to recruit the participants and collect the data.

Finally would like to thank Margaret Gillon-Dowens and the Proactive group for comments on previous versions of this manuscript.

## Footnotes

1 Depending on the theoretical model of reference, these representational dynamics can change a lot. For example, Roelofs (1992) suggests that after lexical-semantic access, an intermediate “lexeme” representation modulates sub-lexical phonological processing. In this framework, we mainly assume that (i) full lexical access effects should emerge earlier in the PMT compared to the WMT and that (ii) later effects imply lower level representations involving information about predictee inflection.

## References

Altmann, G. T., & Kamide, Y. (1999). Incremental interpretation at verbs: Restricting the domain of subsequent reference. Cognition, 73(3), 247–264.

Altmann, G. T., & Mirkovic, J. (2009). Incrementality and prediction in human sentence processing. Cognitive Science, 33(4), 583–609.

Bar, M. (2007). The proactive brain: Using analogies and associations to generate predictions. Trends in Cognitive Sciences, 11(7), 280–289.

Bastos, A. M., Vezoli, J., Bosman, C. A., Schoffelen, J. M., Oostenveld, R., Dowdall, J. R., Fries, P. (2015). Visual areas exert feedforward and feedback influences through distinct frequency channels. Neuron, 85(2), 390–401.

Boersma, Paul (2001). Praat, a system for doing phonetics by computer. Glot International 5:9/10, 341–345.

Caffarra, S., & Barber, H. (2015). Does the ending matter? The role of gender-to ending consistency in sentence reading. Brain Research, 1605, 83–92.

Caffarra, S., Janssen, N., & Barber, H. (2014). Two sides of gender: ERP evidence for the presence of two routes during gender agreement processing. Neuropsychologia, 63, 124–134.

Caramazza, A. (1997). How many levels of processing are there in lexical access? Cognitive Neuropsychology, 14, 177–208.

Chang, F., Dell, G. S., & Bock, K. (2006). Becoming syntactic. Psychological Review, 113(2), 234–272.

Clark, A. (2013). Whatever next? Predictive brains, situated agents, and the future of cognitive science. Behavioral and Brain Sciences, 36(3), 181–204.

Coltheart, M., Rastle, K., Perry, C., Langdon, R., Ziegler, J. (2001). DRC: A dual route cascaded model of visual word recognition and reading aloud. Psychological Review, 108, 204–256.

DeLong, K., Urbach, T. P., & Kutas, M. (2005). Probabilistic word pre-activation during language comprehension inferred from electrical brain activity. Nature Neuroscience, 8(8), 1117–1121

Dehaene, S. Duhamel, J., Hauser, M., and Rizzolatti, G. (2005) From Monkey Brain to Human Brain. A Fyssen Foundation Symposium. MIT Press.

Dell, G. S., & Brown, P. M. (1991). Mechanisms for listener-adaptation in language production: Limiting the role of the “model of the listener”. In D. J. Napoli & J. A. Kegl (Eds.), Bridges between psychology and linguistics: A Swarthmore Festschrift for Lila Gleitman (Vol. 105, pp. 105–129): Psychology Press.

Dikker, S., Rabagliati, H., Farmer, T. A., & Pylkkänen, L. (2010). Early occipital sensitivity to syntactic category is based on form typicality. Psychological Science, 21(5), 629–634.

Duchon, A., Perea, M., Sebastian-Galles, N., Marti, A., Carreiras, M. (2013). EsPal: one-stop shopping for Spanish word properties. Behavioral Research Methods, 45(4), 1246–1258.

Federmeier, K. D. (2007). Thinking ahead: The role and roots of prediction in language comprehension. Psychophysiology, 44, 491–505

Federmeier, K. D., McLennan, D. B., De Ochoa, E., Kutas, M. (2002). The impact of semantic memory organization and sentence context information on spoken language processing by younger and older adults: an ERP study. Psychophysiology, 39, 133–146.

Foucart, A., Martin, C. D., Moreno, E. M., & Costa, A. (2014). Can bilinguals see it coming? Word anticipation in L2 sentence reading. Journal of Experimental Psychology: Learning, Memory, and Cognition, 40(5), 1461–1469.

Friston, K. J. (2005). A theory of cortical responses. Philosophical Transactions of the Royal Society. Biological Sciences, 360(1456), 815–836.

Friston, K. J. (2010). The free-energy principle: a unified brain theory? Nature Reviews Neuroscience, 11(2), 127–138.

Grainger, J., & Holcomb, P. J. (2009). Watching the word go by: On the time-course of component processes in visual word recognition. Language and Linguistic Compass, 3(1), 128–156.

Grainger, J., & Jacobs, A. M. (1996). Orthographic processing in visual word recognition. A multiple readout model. Psychological Review, 103, 518–565.

Gollan, T., & Frost, R. (2001). Two Routes to Grammatical Gender: Evidence from Hebrew. Journal of Psycholinguistic Research, 30, 627–651.

Hawelka, S., Schuster, S., Gagl, B., & Hutzler, F. (2015). On forward inferences of fast and slow readers. An eye movement study. Scientific reports, 5.

Hahne, A. (2001). What’s different in second-language processing? Evidence from event-related brain potentials. Journal of Psycholinguistic Research, 30, 251–266.

Hahne, A., & Friederici, A. D. (2001). Processing a second language: Late learners comprehension mechanisms as revealed by event-related brain potentials. Bilingualism: Language and Cognition, 4(2), 123–141.

Indefrey P., Levelt W. J. M. (2004). The spatial and temporal signatures of word production components. Cognition 92, 101–144.

Ito, A., Martin, A.E., & Nieuwland, M.S. (in press). On prediction of form and meaning in a second language. Journal of Experimental Psychology: Learning, Memory, and Cognition.

Izura, C., Cuetos, F., & Brysbaert, M. (2014). Lextale-Esp: A test to rapidly and efficiently assess the Spanish vocabulary size. Psicologica, 35, 49–66

Jackendoff, R. (2002). Foundations of Language: Brain, Meaning, Grammar, Evolution. Oxford: Oxford University Press. p. 477.

Jaeger, T. F., & Snider, N. E. (2013). Alignment as a consequence of expectation adaptation: syntactic priming is affected by the prime’s prediction error given both prior and recent experience. Cognition, 127(1), 57–83.

Kaan, E. (2014). Predictive sentence processing in L1 and L2. Linguistic Approaches to Bilingualism, 4(2) 257–282.

Kamide, Y., Altmann, G. T., & Haywood, S. L. (2003). The time-course of prediction in incremental sentence processing: Evidence from anticipatory eye movements. Journal of Memory and Language, 49, 133–156

Kutas, M., & Hillyard, S. A. (1984). Brain potentials during reading reflect word expectancy and semantic association. Nature, 307(5947), 161–163.

Kuperberg, G. R., & Jaeger, T. F. (2016). What do we mean by prediction in language comprehension? Language, Cognition and Neuroscience, 31(1), 32–59.

Laka, I. (1996). A Brief Grammar of Euskara, the Basque Language. Euskal Herriko Unibertsitatea: Leioa-Donostia (Spain)

Levelt, W. J. M., Roelofs, A., & Meyer, A. S. (1999). A theory of lexical access in speech production. Behavioral and Brain Sciences, 22, 1–38.

Levy, R. (2008). Expectation-based syntactic comprehension. Cognition, 106(3), 1126–1177

Luck S.J., Hillyard S.A. (1994). Electrophysiological correlates of feature analysis during visual search. Psychophysiology, 31, 291–308.

Luke, S. G., & Christianson, K. (2016). Limits on lexical prediction during reading. Cognitive Psychology, 88, 22–60.

MacDonald, M. (2013). How language production shapes language form and comprehension. Frontiers in Psychology, 4.

Marslen-Wilson W.D. (1987). Functional parallelism in spoken word-recognition. Cognition 25(1–2), 71–102.

Martin, C., Thierry, G., Kuipers, J.-R., Boutonnet, B., Foucart, A., & Costa, A. (2013). Bilinguals reading in their second language do not predict upcoming words as native readers do. Journal of Memory and Language, 69(4), 574–588.

McClelland, J. L., & Rumelhart, D. E. (1981). An interactive activation model of context effects in letter perception: part I: an account of basic findings. Psychological Review, 88, 375.

Molinaro, N., Barber, H.A., Perez, A., Parkkonen, L., & Carreiras, M. (2013). Left fronto-temporal dynamics during agreement processing: Evidence for feature specific computations. Neuroimage, 78, 339–352.

Molinaro, N., Barraza, P., & Carreiras, M. (2013). Long-range neural synchronization supports fast and efficient reading: EEG correlates of processing expected words in sentences. Neuroimage, 72, 120–132.

Molinaro, N., Giannelli, F., Caffarra, S., & Martin C. (2016). The native language tunes prediction processes across multiple languages. Poster presented at the ‘23rd Meeting of the Cognitive Neuroscience Society’, New York, USA.

Molinaro, N., Giannelli, F., Caffarra, S., & Martin C. (under review) Hierarchical levels of representation in language prediction: The influence of first language acquisition in highly proficient bilinguals.

Molnar, M., Lallier, M., & Carreiras, M., (2014). The Amount of Language Exposure Determines Nonlinguistic Tone Grouping Biases in Infants From a Bilingual Environment. Language Learning, 64(s2), 45–64.

Morris, R. K. (2006). Lexical processing and sentence context effects. In M. J. Traxler & M. A. Gernsbacher (Eds.), Handbook of psycholinguistics (2nd ed., pp. 377–402).

Pickering, M. J., & Garrod, S. (2013). An integrated theory of language production and comprehension. Behavioral and Brain Sciences, 36(4), 329–347.

Peirce, JW (2007) PsychoPy-Psychophysics software in Python. Journal of Neuroscience Methods, 162(1–2), 8–13.

R Core Team (2015). R: A language and environment for statistical computing. R Foundation for Statistical Computing. Vienna, Austria.

Rao, R. P., & Ballard, D. H. (1999). Predictive coding in the visual cortex: a functional interpretation of some extra-classical receptive-field effects. Nature Neuroscience, 2(1), 79–87.

Rijk, R.P.G. (2008). Standard Basque, a Progressive Grammar. Cambridge MA: MIT Press.

Real Academia Española. (2009). Nueva gramática de la lengua española. Asociaciòn de academias de la lengua Española.

Roelofs, A., (1992). A spreading-activation theory of lemma retrieval in speaking. Cognition 42, 107–142

Strijkers K, Costa A (2016). On words and brains: linking psycholinguistics with neural dynamics in speech production. Language, Cognition and Neuroscience, 31(4), 524–35.

Wicha, N. Y. Y., Moreno, E. M., & Kutas, M. (2004). Anticipating words and their gender: An event-related brain potential study of semantic integration, gender expectancy, and gender agreement in Spanish sentence reading. Journal of Cognitive Neuroscience, 16(7), 1272–1288

